# Modeling Changes in Probabilistic Reinforcement Learning during Adolescence

**DOI:** 10.1101/2020.12.02.407932

**Authors:** Liyu Xia, Sarah L Master, Maria K Eckstein, Beth Baribault, Ronald E Dahl, Linda Wilbrecht, Anne GE Collins

## Abstract

In the real world, many relationships between events are uncertain and probabilistic. Uncertainty is also likely to be a more common feature of daily experience for youth because they have less experience to draw from than adults. Some studies suggests probabilistic learning may be inefficient in youth compared to adults [1], while others suggest it may be more efficient in youth that are in mid adolescence [2, 3]. Here we used a probabilistic reinforcement learning task to test how youth age 8-17 (N = 187) and adults age 18-30 (N = 110) learn about stable probabilistic contingencies. Performance increased with age through early-twenties, then stabilized. Using hierarchical Bayesian methods to fit computational reinforcement learning models, we show that all participants’ performance was better explained by models in which negative outcomes had minimal to no impact on learning. The performance increase over age was driven by 1) an increase in learning rate (i.e. decrease in integration time horizon); 2) a decrease in noisy/exploratory choices. In mid-adolescence age 13-15, salivary testosterone and learning rate were positively related. We discuss our findings in the context of other studies and hypotheses about adolescent brain development.

**Author summary:** Adolescence is a time of great uncertainty. It is also a critical time for brain development, learning, and decision making in social and educational domains. There are currently contradictory findings about learning in adolescence. We sought to better isolate how learning from stable probabilistic contingencies changes during adolescence with a task that previously showed interesting results in adolescents. We collected a relatively large sample size (297 participants) across a wide age range (8-30), to trace the adolescent developmental trajectory of learning under stable but uncertain conditions. We found that age in our sample was positively associated with higher learning rates and lower choice exploration. Within narrow age bins, we found that higher saliva testosterone levels were associated with higher learning rates in participants age 13-15 years. These findings can help us better isolate the trajectory of maturation of core learning and decision making processes during adolescence.

## Introduction

In the everyday world, perfectly predictable outcomes are rare. Yet, we need to track important events and their relationships to other events and actions. For example, we might want to learn where the best place to obtain food is, or where a potential mate likes to hang out – this might help us decide where to go, expecting a positive outcome to occur frequently, but not always. Our ability to learn about these probabilistic relationships is crucial for our daily life and decision making.

This challenge needs to be met by the developing brain, especially during adolescence [4–9]. Naively, one might assume that the brain simply gets better at this (and possibly all) forms of learning with brain maturation. However, what does *better* mean in this context? Most learning mechanisms are subject to tradeoffs between speed and stability. Fast learning may be suitable for a highly certain environment with deterministic relationships/statistics, but can lead to impulsive behavior in a more uncertain environment with probabilistic relationships/statistics [2, 10]. By contrast, slower and more integrated learning may lead to more robust and stable performance in probabilistic environments. During development, there may be periods where one form of learning is emphasized over the other. Changes could be gradual and monotonic, but there may also be non-monotonic changes, such as inverted U shapes [3, 7, 11]. Peaks in inverted U shapes indicate a time when a phenomenon is at its maximum; such peaks for learning processes during adolescence could accommodate the expected increase in uncertainty in the environment with the transition to independence that occurs at that time of life across species [12].

To study how learning changes across adolescence, we used the theoretical framework of reinforcement learning (RL). Computational RL models assume that we estimate the long-term values of an action in a given state by integrating over time the feedback we receive for choosing this action in this state, through a trial-and-error process [13]. RL has greatly enhanced our understanding of human behavior and the neural processes that underlie learning and decision-making in both certain and uncertain environments [14–18].

RL has also been previously used to probe developmental changes in learning and decision making, including during adolescence [1–5, 11, 19]. While there has been some consensus on certain developmental trends, such as lower decision noise with age [4], in general, developmental results in both how learning behavior changes and in how RL processes (and parameters) change are highly variable and dependent on the specific tasks used [4, 20].

To mitigate the issue introduced by task heterogeneity, we conducted a sequence of four behavioral tasks on the same population of 297 participants across a wide age range (8-30), over-sampling participants age 8-18 to focus on the adolescent period (S1 Fig). The four tasks varied across multiple dimensions (such as deterministic/probabilistic feedback, stable/volatile contingencies, memory load, etc.). The focus of this paper is on one of these tasks, the *Butterfly task* [2], where participants needed to learn probabilistic associations that were stable throughout the task. While we mainly present results from the Butterfly task, we also discuss comparisons and relationships with two other tasks in this sequence of four tasks [3, 11].

The Butterfly task tests participants’ ability to learn probabilistic associations between four butterflies and two possible preferred flowers from reward feedback. While this task has been used in developmental studies before [2], it included other features that could affect the learning process (novel images unrelated to the butterflies), and was run on a relatively small sample. Here, we sought to investigate in a much larger sample how participants’ performance in a probabilistic reinforcement learning task changed across adolescence, in the absence of the novel images.

While the previous study found better performance in adolescents than adults [2], we found that performance increased through early twenties, then stabilized, peaking at around 25 years. We used hierarchical Bayesian methods to fit computational RL models to the trial-by-trial data (see Hierarchical model fitting) and examined how subjects integrated information across trials and made decisions. Increases in performance with age were explained by an increase in learning from rewarded outcomes and a decrease in exploration. These findings are largely consistent with a general picture emerging from studies of learning and decision making across development [2, 4, 11]. However, we also discuss some notable differences [2, 3].

## Materials and methods

### Participants

All procedures were approved by the Committee for the Protection of Human Subjects (CPHS number, community participants: 2016-06-8925; student participants: 2016-01-8280) at the University of California, Berkeley (UCB). A total of 297 participants completed the task: 187 children and adolescents (age 8-18) from the community, 55 UCB undergraduate students (age 18-25), and 55 adults (age 25-30) from the community. Participants under 18 years old and their guardians provided their informed assent or written permission; participants over 18 provided informed written consent themselves.

We assessed pubertal development for children and adolescents through saliva samples and the pubertal development questionnaire, from which we calculated Puberty Development Score (PDS, [21]) and testosterone levels (T1, see Saliva collection and testosterone testing).

Community subjects were compensated with a $25 Amazon gift card for completing the experimental session, and an additional $25 for completing optional take-home saliva samples; undergraduate participants received course credit for participation. All participants were pre-screened for the absence of present or past psychological and neurological disorders.

### Saliva collection and testosterone testing

In addition to self-report measures of pubertal development, we also collected saliva from each of our subjects to quantify salivary testosterone, following methods reported in [11]. Testosterone is a reliable measure of pubertal status in boys and girls and is associated with changes in brain and cognition in adolescence [22, 23]. Subjects refrained from eating, drinking, or chewing anything at least an hour before saliva collection. Subjects were asked to rinse their mouth out with water approximately 15 minutes into the session. At least one hour into the testing session, they were asked to provide 1.8 mL of saliva through a plastic straw into a 2 mL tube. Subjects were instructed to limit air bubbles in the sample by passively drooling into the tube, not spitting. Subjects were allotted 15 minutes to provide the sample. After the subjects provided 1.8 mL of saliva, or 15 minutes had passed, the sample was immediately stored in a −20°F freezer. The date and time were noted by the experimenter. The subjects then filled out a questionnaire of information which might affect the hormone concentrations measured in the sample (i.e. whether the subjects had recently exercised).

Salivary testosterone was quantified using the Salimetrics Salivatory Testosterone ELISA (cat. no. 1-2402, Bethesda, MA). Intra- and inter-assay variability for testosterone were 3.9% and 3.4%, respectively. Samples below the detectable range of the assay were assigned a value of 5 pg/mL, 1 pg below the lowest detectable value. Final testosterone sample concentration data were cleaned with a method developed in [24]. There were no subjects with any samples above the detectable range. Within subjects aged 8 to 17 only, outliers greater than 3 standard deviations above the group mean were fixed to that value, then incremented in values of +0.01 to retain the ordinality of the outliers.

### Experimental design

This task was the third in a sequence of four tasks that participants completed in the experimental session [11]. The task was a contextual 2-armed bandit task with binary feedback: there were 4 stimuli (blue, purple, red, and yellow butterflies) and 2 bandits (pink and white flowers). Participants needed to figure out the preferred flower for each of the four butterflies through trial and error. Each butterfly had a preferred flower, which remained fixed throughout the experiment.

On each trial (Fig 1A), participants were presented one butterfly and needed to choose a flower within 7s using a video game controller. They were instructed to respond as quickly as possible. The chosen flower would stay on the screen for 1s. If participants correctly chose the preferred flower, they would receive positive feedback (*Win!*) with 80% chance; however, 20% of the time the other flower would be the rewarding one, resulting in negative feedback (*Lose!*). The feedback stayed on the screen for 2s. There were 30 trials for each butterfly, resulting in a total of 120 trials.

**Fig 1.**
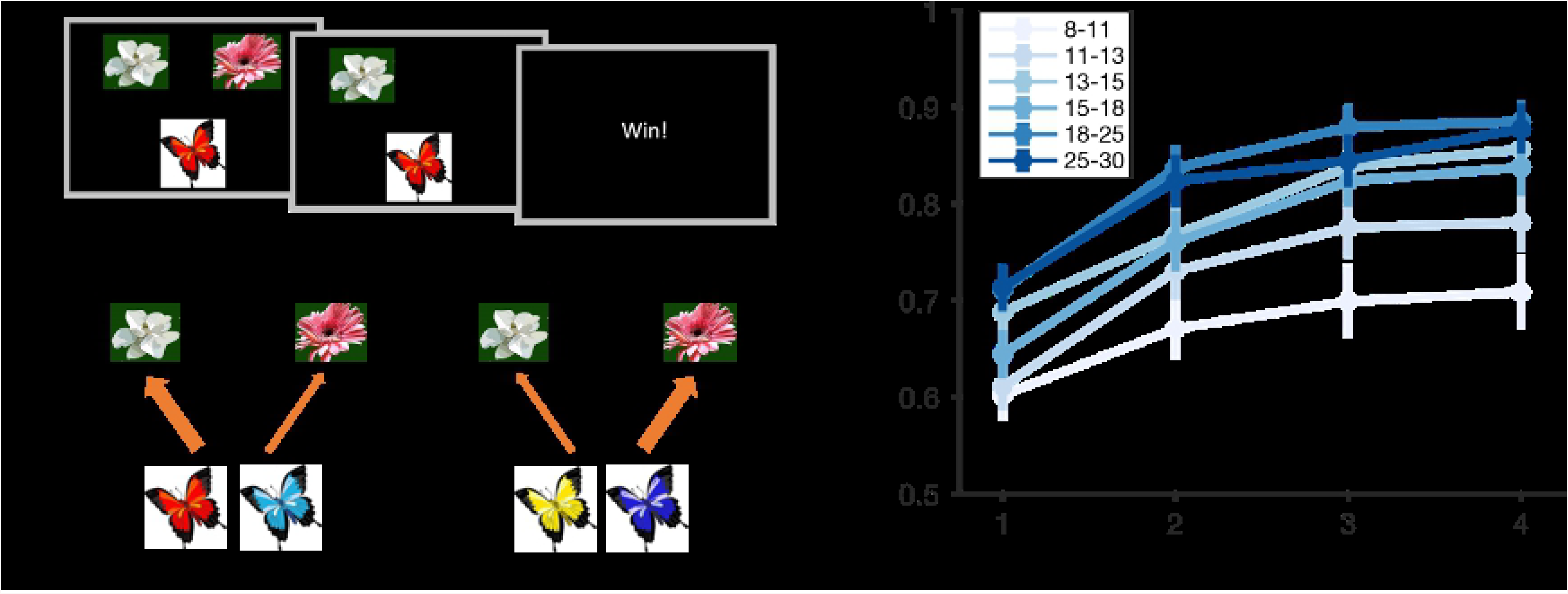
Experimental design and overall performance. (A) On each trial, participants needed to choose the butterfly’s preferred flower. Each butterfly’s preferred flower stayed the same throughout the experiment. If participants correctly picked the butterfly’s preferred flower, they observed a *Win!* feedback with 80% chance, and *Lose!* otherwise. The other choice delivered positive feedback only with 20% chance. (B) Average probability of a correct choice over four 30-trial learning blocks. Learning curves showed all age groups learned the task, and that performance generally improved with age group.

The butterfly-flower mapping, position of flowers, sequence of butterflies and the probabilistic feedback were pre-randomized and counterbalanced across participants.

### Exclusion criteria

We excluded participants who were overall more likely to change their choice of flower for a given butterfly (switch) than repeat it after receiving positive feedback, which suggested that they either did not understand the task or were not engaged in it. 20 participants under 18 and 1 undergraduate participant were excluded due to this criterion.

To further conservatively identify participants who were not engaged in the task, we excluded participants who had worse than chance performance and satisfied one of the following “low data quality” criteria: (1) high proportion of trials where the participant picked the same choice as the previous trial, (2) high proportion of trials where the participant changed their choice, (3) presence of too long sequences of trials where the participant kept choosing the same flower, and (4) high proportion of missing trials. All of those criteria were determined by elbow points (see Exclusion criteria details), and indicated a lack of reactivity to the task’s inputs. After applying these conjunctive criteria, the total number of participants for later analysis was 264, with 157 participants under 18 (see S1 Table for breakdown of exclusion by age). All results presented hold with weaker exclusion criteria.

### Model-independent analysis

For each participant in each trial, we recorded whether they chose the butterfly’s preferred flower (*correct* choice or not), and whether they received reward or not (win vs. lose), which was different due to the probabilistic nature of the task. As an aggregate measure of performance, we computed average accuracy within each of the four 30-trial learning blocks for each participant. We also computed median and standard deviation of reaction time within each learning block. We ran (linear and quadratic) regression to assess whether those behavioral metrics changed with age and pubertal measures.

We also calculated the proportion of trials (*p*) among all 120 trials where participants correctly chose each butterfly’s preferred flower as an overall performance measure. Because this proportion was not normally distributed across participants (Kolmogorov–Smirnov test, *p* = 0.003), we instead used log odds 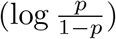 for all later statistical tests. The log odds were normally distributed (Kolmogorov–Smirnov test, *p* = 0.26).

We used the median reaction time for each participant as a speed measure. Because reaction time was not normally distributed across participants (Kolmogorov–Smirnov test, *p* = 0.02), for all later statistical tests, we used log-transformed reaction time, which was normally distributed (Kolmogorov–Smirnov test, *p* = 0.8).

To visualize age effects (Fig 1, 2), we broke participants under age 18 into 4 equal-size groups within each sex respectively, and then combined both sexes. The boundaries for the four age groups were approximately 8-11, 11-13, 13-15, and 15-18. Together with two age groups above 18 (18-25 for students and 25-30 for community subjects), we had a total of six age groups.

**Fig 2.**
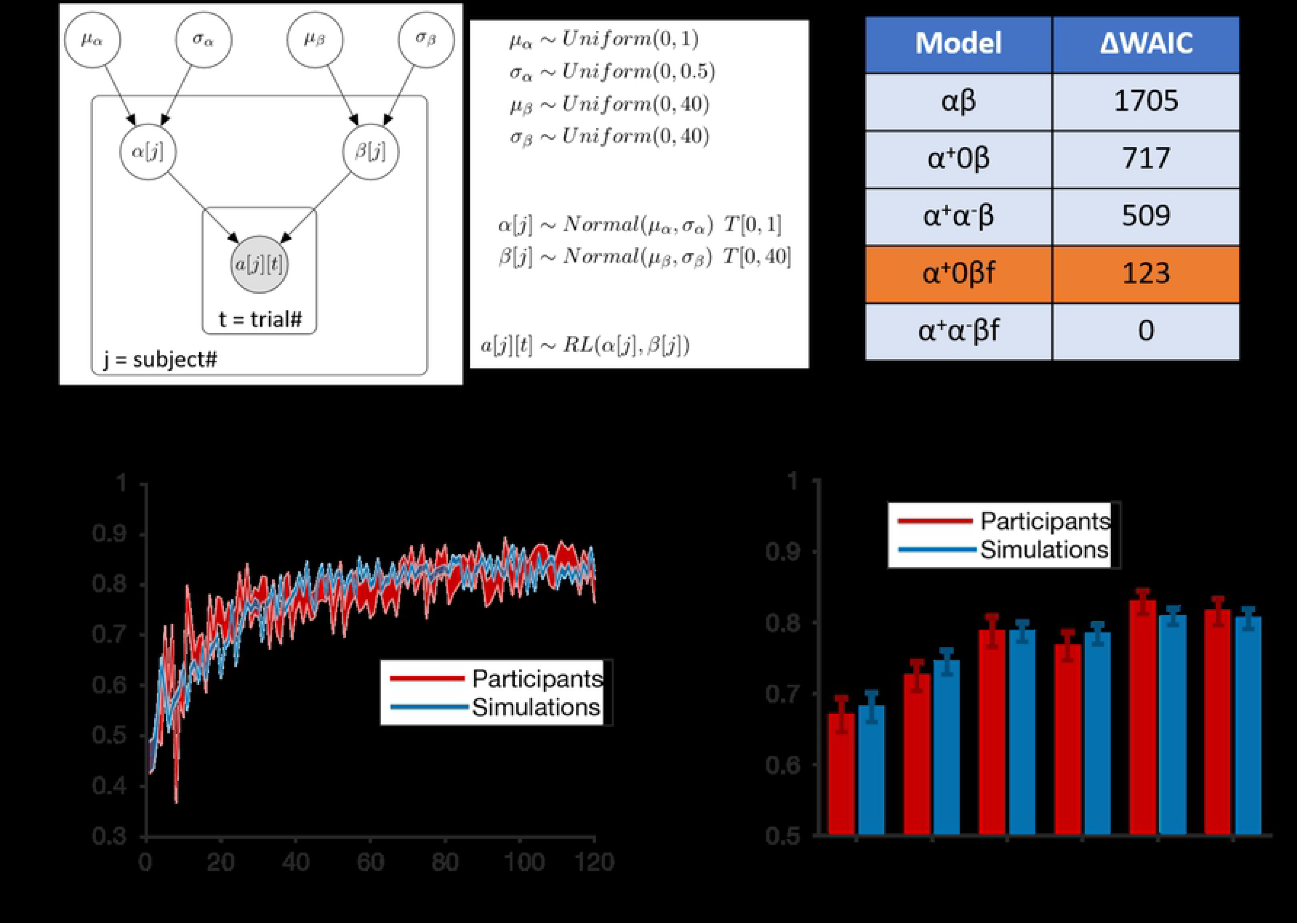
Hierarchical Bayesian modeling and model comparison. (A) We illustrate the hierarchical model design with the *αβ* model as an example, We assumed that parameters of individual participants came from the same group level distribution, which is truncated normal parametrized by the group-level mean and standard deviation (*μ*_*α*_ and *σ*_*α*_ for *α*; *μ*_*β*_ and *σ*_*β*_ for *β*). The group-level mean and standard deviation followed weakly informative priors (uniform and bounded). The parameters for each individual participant are used in the likelihood of each action on every trial based on the *αβ* model. *T* [*m, n*] indicates truncation of distribution (e.g. the learning rate, *α*, is bounded by [0, 1]). The filled circle represented observed variable (in this case, participants’ choices on each trial); unfilled circles represented latent variables (in this case, group and individual model parameters). (B) We calculated WAIC for model comparison. While the *α*^+^*α*^−^*βf* model had the lowest (i.e. best) WAIC score, the *α*^*−*^ parameter was not recoverable and showed signs of overfitting (see Model comparison extended). We thus focus on the *α*^+^0*βf* model. (C-D) We used fitted parameters from the *α*^+^0*βf* model to generate simulated trajectories. The simulated performance captures the average learning curve throughout the experiment (C), and replicates the age effect (D).

Going beyond aggregate measures across trials, we also ran a mixed effect logistic regression to predict participants’ choices on a trial-by-trial basis and tested how previous reward history and delay affected learning and decision making. Specifically, for each trial, we defined the *reward history*, *r*, as the number of trials that participants had previously received a *Win!* feedback for the current butterfly; we also defined *delay*, *d*, as the number of intervening trials since the last time the participant encountered the same butterfly and got rewarded. We then used the lme4 package in R to test *p*(*correct*) = *logit*(1 + *r* + *d* + (1 + *r* + *d | sub*)), where *sub* represented random effects of individual participants respectively. All regressors were z-scored. We analyzed whether the random effects varied with age using linear and quadratic regressions.

### Computational models

We used computational RL modeling to obtain a more quantitative and mechanistic understanding of participants’ trial-by-trial learning. We applied five variants of RL models, then used the parameter estimates of the best fitting model as the basis for inference.

#### Classic RL (*αβ*)

Our simplest RL model was the *αβ* model, with just 2 free parameters, *α* (learning rate) and *β* (inverse temperature). The *αβ* model used Q-learning to compute *Q*(*b, a*), as the expected value of choosing flower *a* for butterfly *b*. On trial *t*, the probability of choosing *a* was computed by transforming the Q-value with a softmax function:

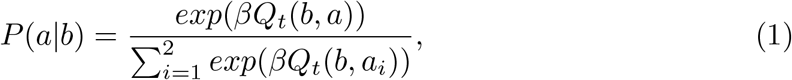

where *Q*_*t*_(*b, a*) was the Q-value until trial *t*. After observing reward *r*_*t*_ (0 for “Lose!” or 1 for “Win!”), the Q-value *Q*(*b, a*) was updated through the classic delta rule:

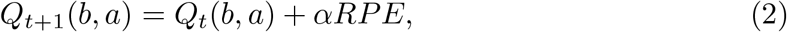

where *RPE* = *r*_*t*_ − *Q*_*t*_(*b, a*) is the reward prediction error. We initialized all Q-values to the uninformative value of 0.5 (the average of positive and negative feedback) for this model and all other models under consideration.

#### RL with asymmetric learning rates (*α*^+^*α^−^β*)

The *α*^+^*α*^−^*β* model differed from the *αβ* model by using two distinct learning rate parameters, *α*^+^ and *α*^−^. This allowed the model to have different sensitivity to positive and negative RPE [25]. Specifically, in Eq 2, *α*^+^ was used when *RPE* > 0, and *α^−^* otherwise.

#### Asymmetric RL with *α*^−^ = 0 (*α*^+^0*β*)

The *α*^+^0*β* model was the same as the *α*^+^*α*^−^*β* model, except that the *α*^−^ parameter was set to 0. This change made the model insensitive to negative feedback. We included this model because of the observation that the fitted values of the *α^−^* parameter from the *α*^+^*α*^−^*β* model were very small and not recoverable (see Model comparison extended).

#### RL with forgetting (*α*^+^0*βf*)

The *α*^+^0*βf* model builds upon the *α*^+^0*β* model by including an additional forgetting parameter, *f*. On each trial, after applying the delta learning rule Eq 2, Q-values decay toward the uninformative starting value of 0.5, implementing a forgetting process:

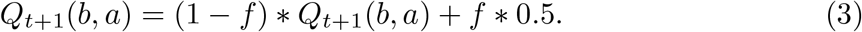

Eq 3 is implemented for all butterfly-flower pairs except the butterfly and the selected flower on the current trial.

#### RL with asymmetric learning rates and forgetting (*α*^+^*α^−^βf*)

For completeness, we also included the *α*^+^*α^−^βf* model, which has both asymmetric learning rates for positive and negative feedback and the forgetting parameter.

### Hierarchical model fitting

We fitted all five RL models using hierarchical Bayesian methods [26] jointly to all participants, instead of to each participant independently. To illustrate the hierarchical model design (Fig 2A), we use the simplest model, *αβ*, as an example. We began by specifying weakly informative priors for the mean and standard deviation of the group-level learning rate (*μ*_*α*_ and *σ*_*α*_) and the group-level inverse temperature (*μ_β_* and *σ_β_*). We assumed that these group-level parameters were all uniformly distributed over the natural ranges of the parameters (for example, we truncated *μ*_*α*_ at 0 and 1, since we know the learning rate parameter is between 0 and 1). We then assumed that the parameters for each participant were drawn from a prior distribution defined by the group-level parameters: for example, *α*[*j*] for participant *j* was drawn from a normal distribution *Normal*(*μ*_α_, *σ*_*α*_) truncated at 0 and 1. Individual participants’ parameters were then used in the likelihood of each participant’s actions on each trial (*a*[*j*][*t*]) according to the *αβ* model, where *j* and *t* indicate participant number and trial number, respectively.

The hierarchical model made the likelihood intractable [27], but it can be well approximated by sampling. We used No-U-Turn sampling, a state-of-the-art Markov Chain Monte Carlo (MCMC) algorithm implemented in the probabilistic programming language Stan [28], to sample from the joint posterior distribution of model parameters for all participants. Compared to the classic participant-wise maximum likelihood estimation approach, hierarchical model fitting with MCMC provides more stable point estimates for individual participants and allows natural inference of effects on parameters at the group level [29].

For each model, we ran 4 MCMC chains, with each chain generating 4000 samples (after 1000 warmup samples), resulting in 16000 samples per model for later inference. We assessed convergence for all models using the *matstanlib* library [30]. In particular, we ensured that 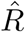 statistics for all free parameters were below 1.05; that the effective sample sizes (ESS) for all free parameters were more than 25 × the number of chains; and that samples generally were not the result of divergent transitions. Note that these criteria are more stringent than the standard criteria for convergence of 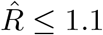 and ESS > 5× number of chains as per [26].

Hierarchical Bayesian modeling also provides a natural way to test for potential age effects on model parameters. Specifically, we incorporated the regression model using age to predict model parameters into the original graphical model (Fig 2A), and directly sampled regression coefficients for age jointly with other model parameters.

Same as before, we assumed that the model parameters for individual participants followed a truncated normal distribution with a group-level standard deviation, but now we replace the prior on the group-level mean with a regression statement with respect to age. For example, to probe linear effects of age on *α*, we assumed that the parameter *α*[*j*], used to compute the likelihood of participant *j*’s choices, followed *Normal*(*α*_*intercept*_ + *α*_*linear*_ * *age*[*j*], *σ*_*α*_), *T* [0, 1], where *age*[*j*] was the z-scored age of participant *j*, and *α*_*intercept*_, *α*_*linear*_ were regression coefficients for which we set weakly informative priors. To probe quadratic effects, we just need to further include *α_quadratic_* * *age*[*j*]^2^.

To test for effects of age on the model parameters, we examined whether the posterior distribution of all 16000 samples for the linear (*α*_*linear*_) and quadratic (*α*_*quadratic*_) regression coefficients were significantly different from 0.

## Results

### Overall performance

To assess learning progress and potential age effects, we first calculated the proportion of correct trials within each of the four 30-trial learning blocks for six age groups (Fig 1B). As indicated in methods, we grouped all participants under 18 into 4 equal-sized bins (N = 39, 39, 39, 40). The other two groups were undergraduate participants (age 18-25, N = 53) and adult community participants (age 25-30, N = 54).

All age groups exhibited learning over the course of the experiment. Specifically, we found a significant main effect of age group and block on participants’ performance (2-way mixed-effects ANOVA, age group: *F* (5, 255) = 8.5*, p* < 0.0001; block: *F* (3, 765) = 136, *p* < 0.0001). There was no interaction between age group and block (2-way mixed-effects ANOVA: *F* (15, 765) = 1, *p* = 0.49). This shows that participant’s performance improved as the experiment progressed, and older participants generally outperformed younger participants.

To further characterize the effect of age on overall performance, we computed the proportion of correct trials over all 120 trials. We found that the overall performance of 13-18 year-olds (top-two quartiles) was significantly higher than of 8-13 year-olds (bottom two quartiles; unpaired t-test, *t*(1, 155) = 3.5, *p* = 0.0001), and significantly lower than of 18-25 year-olds (unpaired t-test, *t*(1, 130) = 2.5, *p* = 0.01). However, there was no significant difference (unpaired t-test, *t*(1, 105) = 0.2, *p* = 0.8) between the performance of 25-30 year-olds and 18-25 year-olds (Fig 1B, Fig 2D).

To examine the continuous relationship between participants’ performance and age, we ran a regression analysis using age to predict performance (Fig 3A). We found that including a quadratic term of age improved fit in terms of the Akaike Information Criterion (AIC [31]; AIC(linear) = 671; AIC(quadratic) = 666). The regression analysis revealed linear and quadratic effects of age on performance (linear: *β*_*age*_ = 0.05, 95% CI = [0.03, 0.07]; quadratic: 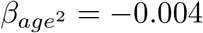, 95% CI = [−0.007, −0.001]). This indicates an inverse-U shape performance curve, with maximal performance around age 25, confirming the previous group analysis. There was no effect of sex or interaction with age (multiple linear regression, both *p*’s > 0.45).

**Fig 3.**
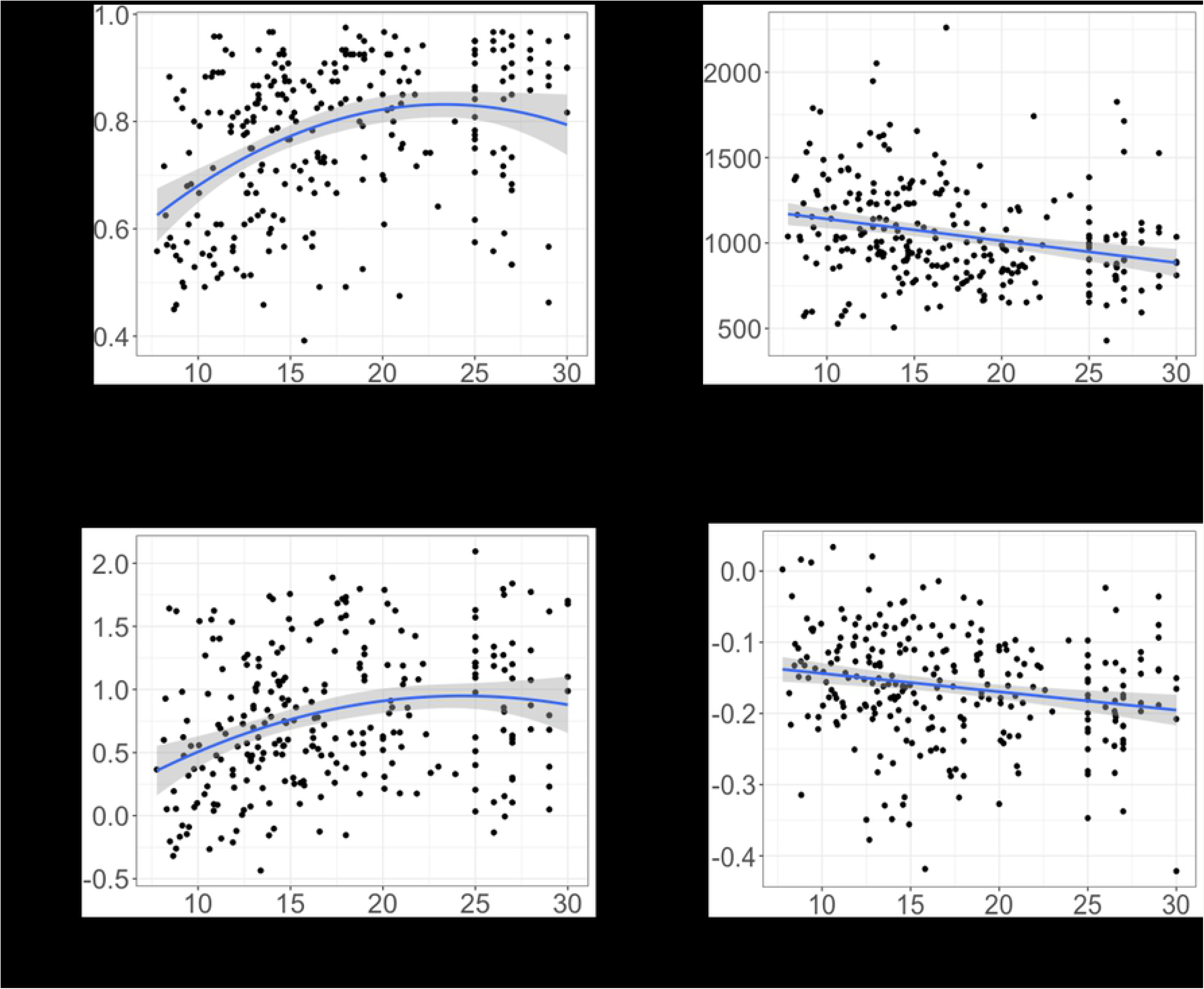
Age effects on participants’ behavior. Scatter plot of age (x-axis) and (A) probability of choosing the correct response, (B) median reaction time, (C) random effect for reward history, and (D) random effect of delay. Each black dot represents one participant. The blue curve represents linear/quadratic regression line. There was no effect of sex in any analysis. Shaded region represents 95% confidence interval.

### Reaction time

We also computed the median (Fig 3B) and standard deviation of reaction time for each participant. We found a linear effect of age on median reaction time (*β*_*age*_ = −0.01, 95% CI = [−0.02, −0.006]). This suggests that participants reacted faster with age, confirming previous results [11]. Adding a quadratic term did not improve fit. We also found a linear effect of age on the standard deviation of reaction time (linear regression: *β*_*age*_ = −0.02, 95% CI = [−0.03, −0.01]); adding a quadratic term provided a better fit (AIC(linear) = 328, AIC(quadratic) = 311, 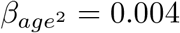, 95% CI = [0.002, 0.005]). This indicates that the variability in reaction time decreased with age, and this decrease itself slowed down with age, consistent with previous findings [11, 32]. There was no significant effect of sex on the median reaction time (unpaired t-test, median: *t*(1, 262) = 0.4*, p* = 0.7), but female participants had a significantly smaller standard deviation than male participants (unpaired t-test: *t*(1, 262) = 2.72*, p* = 0.0085).

These results indicate better performance and faster responses in older participants, ruling out speed-accuracy tradeoffs. Both age group (Fig 1B) and continuous age (Fig 3A) analyses revealed an inverse-U shape with performance, suggesting that the age effect slowed down in early adulthood, and might even invert.

### Mixed-effect logistic regression

To better probe trial-by-trial learning dynamics, we used reward history and delay to predict the probability of a correct choice on each trial in a mixed-effect logistic regression. We found significant fixed effects of reward history and delay (*β*_*r*_ = 0.8, *β*_*d*_ = −0.17, both *p*’s < 0.0001). This suggests that participants were more likely to pick the preferred flower as they received more reward feedback for the butterfly (reinforcement learning effect), and encountered the butterfly more recently (forgetting effect).

We found linear and quadratic effects (linear: *β*_*age*_ = 0.02, 95% CI = [0.01, 0.03]; quadratic: 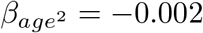, 95% CI = [−0.004, 0.0003]) of age on the random effect of reward history (Fig 3C). This suggests that participants’ sensitivity to reward increased with age and slowed down in adulthood, similar to the trend for overall performance (Fig 3A). Somewhat surprisingly, we also found that participants became more sensitive to delay with age, shown by the linear effect (*β*_*age*_ = −0.003, 95% CI = [−0.004, −0.001]) of age on the random effect of delay (Fig 3D). This could be interpreted in one of two ways: 1) more forgetting over age, or 2) more reliance on decay-sensitive working memory for learning over age.

### Computational modeling

We used computational modeling and model comparison to obtain a mechanistic understanding of participants’ trial-by-trial learning and decision making. We fitted all participants jointly using hierarchical Bayesian modeling [26] combined with sampling [28] for approximating the likelihood function (see Hierarchical model fitting).

#### Model comparison

We used WAIC to compare the relative fit of models at the population level [33], an information criterion that penalizes model complexity appropriately for hierarchical Bayesian models. WAIC is fully Bayesian and invariant to reparametrization. Smaller WAIC indicates a better fit to the data, controlling for complexity. The *α*^+^*α*^−^*βf* model with asymmetric learning rates and the forgetting parameter had the lowest (best) WAIC score (Fig 2B). However, a generate and recover procedure [34] showed (see Model comparison extended) that the *α*^−^ parameter was not recoverable in the *α*^+^*α^−^βf* model, and therefore unsuitable to use as the basis for inference.

Consequently, we focus on the model with the next best WAIC score, *α*^+^0*βf*, which could be successfully recovered from, for further analysis. Note that conclusions for the *α*^+^, *β*, and *f* parameters remain the same if we use the *α*^+^*α*^−^*βf* instead. Furthermore, the *α*^−^ parameters were generally very small for the fitted *α*^+^*α*^−^*βf* model (S2 Fig A). This suggests that participants were learning either very little from negative feedback or not at all. Moreover, the *α*^+^0*βf* model resulted in better model validation (see Model comparison extended), suggesting that *α*^+^*α*^−^*βf* at the population level might be overfitting.

We validated the best-fitting model, *α*^+^0*βf*, by simulating synthetic choice trajectories from fitted parameters (i.e., by generating posterior predictive distributions; Fig 2CD) [35]. Model simulations captured the average learning curve throughout the entire experiment (Fig 2C) and age effects on overall performance (inverse U shape of performance against age groups; Fig 2D).

#### Age differences in model parameters

With the winning model *α*^+^0*βf*, we next asked which computational processes drove the changes in performance over age by looking at how model parameters changed with age. We adapted hierarchical Bayesian modeling to probe effects of age on model parameters. Specifically, we incorporated the regression of age as a predictor of model parameters into the hierarchical Bayesian model (Fig 2A), and directly sampled regression coefficients for age jointly with other model parameters (see Hierarchical model fitting).

To test for effects of age on the model parameters, we examined whether the 95% credible interval (CI) of the posterior samples for each the linear and quadratic regression coefficients did or did not include 0, where 0 indicates no effect (Fig 4). We found linear and quadratic effects of age on *α*^+^ (linear coefficient 95% CI = [0.05, 0.11]; quadratic coefficient 95% CI = [−0.1, −0.03]) and *β* (linear CI = [1.6, 3.4]; quadratic CI = [−1.9, −0.3]). The trajectory of quadratic change over age for *α*^+^ and *β* closely mimicked that for overall performance (Fig 3A). We also found weak linear, but not quadratic effects of age on *f* (linear CI = [−0.04, 0.001]; quadratic CI = [−0.01, 0.02]), with the forgetting parameter decreasing over age.

**Fig 4.**
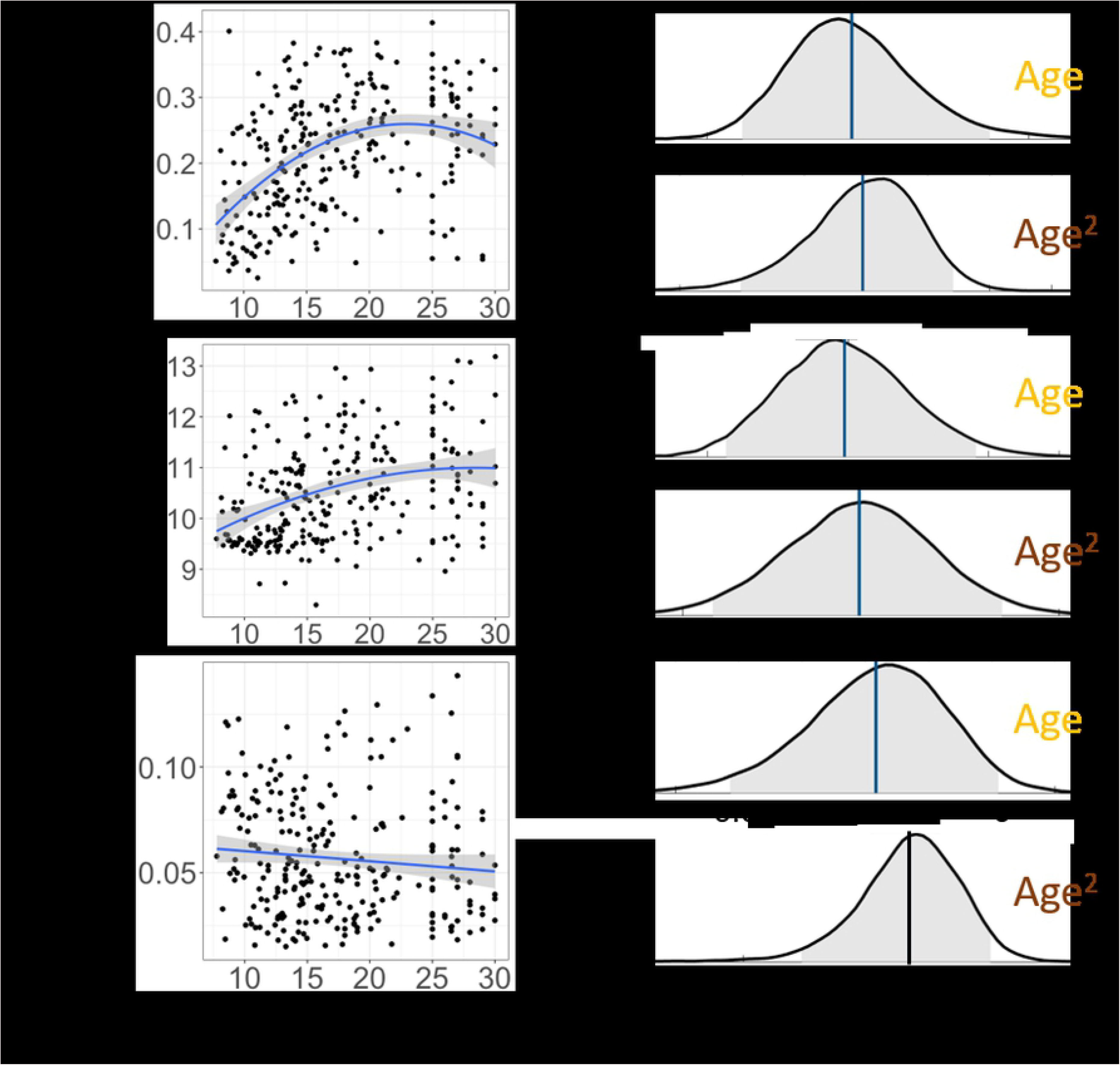
Age effects on model parameters. We directly incorporated age-related parameters into MCMC sampling to test within the hierarchical Bayesian modeling framework whether age had a linear or quadratic effect on all three model parameters: *α*^+^ (A), *β* (B), *f* (C). Left panel: individual parameters from the original *α*^+^0*βf* model plotted against age. For visualization, we included a regression line; the shaded region indicates 95% CI. Right: distribution of posterior samples for linear (top, yellow) and quadratic (bottom, brown) regression coefficients. The vertical line represents the mean of all samples, with blue indicating an effect being present (i.e., 95% CI not including 0), and black indicating no effect. Shaded region shows 95% confidence interval.

### Pubertal effects

To study whether puberty also affected participants’ learning and decision making, we used pubertal measures (pubertal development score PDS and testosterone level T1) to predict fitted model parameters. We found that both PDS and T1 had a linear effect on *α*^+^ (linear regression. PDS: *β*_*PDS*_ = 0.02, 95% CI = [0.01, 0.04]; T1: *β*_*T*1_ = 0.02, 95% CI = [0.01, 0.04]), and no effect on the forgetting parameter *f* (linear regression. Both *p*’s > 0.5). There was a linear effect of PDS, but not T1, on *β* (linear regression. PDS: *β*_*PDS*_ = 0.15, 95% CI = [0.003, 0.29]; T1: *β*_*T*1_ = 0.13, 95% CI = [−0.01, 0.27]). However, the effects of PDS and T1 on *α*^+^ and *β* disappeared when adding age into the regression (multiple linear regression, all *p*’s > 0.55), while age remained the only significant predictor (multiple linear regression, *p*’s < 0.027).

To further explore the effect of PDS and T1 on model parameters while controlling for age, we performed the same regression using PDS or T1 to predict model parameters within each of the four age groups under 18 (S4 Fig, S5 Fig). We found that only within the third age group (age 13-15), there was a positive effect of T1 on *α*^+^ (linear regression: *β*_*T*1_ = 0.05, 95% CI = [0.01, 0.08], *p* = 0.005). This remained significant when correcting for multiple comparisons (two parameters by four groups). This T1 effect remained when controlling for age in the regression (multiple linear regression: *β*_*T*1_ = 0.05, *p*(*T* 1) = 0.006, *p*(*age*) = 0.95). T1 did not provide additional explanatory power for *β* and *f* parameters (see Pubertal effects extended).

## Discussion

How do humans learn to make choices when the outcome is uncertain? To learn probabilistic contingencies, humans need to integrate information over multiple trials to avoid overreacting to noise in the environment. But to learn efficiently, they also need to pay attention to recent information. Here, we investigated how humans trade off these constraints across development, what the underlying computational mechanisms that support such learning are, and how they change during adolescence.

At the population level, computational model comparison (Fig 2B) suggested that two mechanisms modulated learning of probabilistic contingencies. First, participants did not treat positive and negative feedback identically; rather, they had a strong bias to learn more from positive, and little to none from negative feedback. This asymmetry has been widely observed in previous studies [1, 11, 36], potentially due to differential mechanisms integrating positive and negative feedback [37]. Second, we found that learning was better explained by including a forgetting mechanism: more intervening trials between two iterations of a choice decreased the strength of past information [11].

Consistent with the age effects observed in previous work using tasks with probabilistic [3] and deterministic [11] feedback, our behavioral and modeling results suggest that learning in a stable probabilistic task environment changed markedly from childhood to adulthood. In particular, we found that overall performance increased with age, stabilising in early adulthood. This behavioral pattern was mirrored by the learning rate parameter (*α*^+^) as well as inverse temperature (*β*), a parameter indicating a decrease in noise or exploration in choice.

Our observations that learning rate *α*^+^ and inverse temperature *β* increase with development are generally consistent with previous work using the deterministic learning task *RLWM*, tested in the same participants as shown here [11], and a probabilistic task with same the same overall task structure as the Butterfly task, but different feedback methods [2]. However, we did not find higher performance in adolescents than adults, as had been observed in this previous Butterfly task study [2] (Fig 3A). Even when using the same age bins as [2], which limited our subject numbers to N = 89 13-18 year old adolescents and N = 83 20-30 year old adults, we instead found that the performance in 20-30-year-olds was significantly higher than 13-18-year-olds (unpaired t-test, *t*(170) = 2.2, *p* = 0.03).

The finding in [2] was interpreted as “an upside” to slower learning that led to more robust integration over time of information, and thus higher overall performance under uncertainty at younger ages. Indeed, lower learning rates can be more optimal in probabilistic tasks than higher learning rates. However, the relationship between learning rates and performance when learning probabilistic contingencies is complex and non-monotonic: it follows an inverse U-shape, as very low learning rates lead to integrating information too slowly, but high learning rates lead to being too susceptible to noisy feedback [34]. Furthermore, the inverse U-shape itself is dependent on the degree of exploration [2, 4, 34]. Learning rates were smaller in our study compared to [2]: the group level mean for *α*^+^ in our sample was 0.18, whereas in [2], the mean was around 0.3 and 0.6 for adolescents and adults respectively (Fig 2B in [2]). In higher ranges of learning rates [2], an increase in learning rate could result in a decrease in performance (right side of the inverse U-shape), while in our lower range, it could lead to an increase in performance (left side of the inverse U-shape). Thus, the two studies are consistent in identifying an increase in learning rate with age, but over a different range of learning rate values (0.3 vs. 0.6), leading to opposite effects on performance.

Moreover, we modeled learning from positive and negative feedback asymmetrically [4], as opposed to the symmetric learning rate in [2]. In particular, our winning model *α*^+^0*βf* did not learn from negative feedback at all. A high *α*^−^ can also result in worse asymptotic performance in the Butterfly task, resulting in more switching from the preferred flower.

Therefore, while we found a similar trend as in [2] that learning rates increased with age (Fig 4A), our learning rate values were much smaller, and the resulting trend in overall performance was different. Note that this difference in the range of learning rates could be a result of differences in the task specifics (our experiment did not have a memory retrieval aspect with novel images or brain imaging; our task was also the third in a sequence of four tasks). Differences in performance could also stem from differences in socioeconomic status and education level between the groups recruited to each study. For example, it is possible that the peak in performance around age 25 in our sample (Fig 1A) might be driven by the fact that our 18-25 year-olds were undergraduate students, who may have a different education level than the 25-30 year-old community participants in our study or the adults sampled in [2].

Nevertheless, our results support other previous developmental findings. In particular, we also found a decrease in exploration with age [11, 38], and an increase in learning rate previously observed in both deterministic [11] and probabilistic learning tasks [3]. Note that other studies have observed a decrease in learning rates (e.g., 1-learning-rate models: [19, 39, 40]; models with asymmetric learning rates: [1, 36, 41]) or no change [42, 43]. These differences are potentially due to different task structures, samples, and modeling choices. For a more comprehensive review, see [4]).

While we found that performance increased during adolescence and saturated in early adulthood in this stable probabilistic learning task, a probabilistic switching task in the same subjects [3] found a pronounced inverse U shape in overall performance, which peaked at age 13-15. We conclude that this difference in age of peak performance in these two tasks stems from the reliance or lack of reliance on negative outcomes. The probabilistic switching task used in [3] was volatile, and success required learning from negative rewards to follow a switch. In the Butterfly task where the associations were stable throughout the experiment, participants were learning little to nothing from negative feedback. This suggests that even with the same population of participants in two probabilistic tasks, task stability / volatility greatly changed participants’ use of neural systems and behavioral strategies. For this population of subjects, the volatile condition in [3] gave the 13-15 year old adolescents an edge over adults, while the stable condition in the Butterfly Task gave young adults an edge over adolescents.

While we surprisingly found that random effects of delay on performance became more pronounced with age (Fig 3D), we found forgetting became weaker (Fig 4C). One possible interpretation for this apparent contradiction may relate to two simultaneous changes. First, adults may rely more on working memory processes [16] for probabilistic tasks [44], which manifests in strengthened effect of delay. However, the decay of these memory processes may also decrease with age [11], which may be captured here by the decrease in the forgetting parameter. Thus, younger participants might show a weaker effect of delay not because their memory system is forgetting less (it is forgetting more), but because they use their working memory system less in this task, and instead rely more on slower but more robust learning systems.

While we found that pubertal measures did not explain much additional variance compared to age in model parameters (see Pubertal effects extended), we found that testosterone level T1 had a significant positive effect on *α*^+^ within the third age period of 13-15 years. This coincides with the result from [7] that showed a positive relationship between testosterone level and nucleus accumbens activity. These data combined are consistent with a putative link between testosterone, nucleus accumbens activity, and learning rate in mid adolescence.

Another way of interpreting the relationship between testosterone levels and learning rate in age 13-15 can potentially be from the perspective of social learning. One of the most important sources of uncertainty during adolescence comes from social experiences (the rapid changes of social roles and contexts). A previous study [45] indeed showed that testosterone level affects social learning in adolescents. Even though our study did not directly address social learning, the relationship between testosterone and learning might translate to the current paradigm.

## Conclusion

In conclusion, we sought to examine the development of learning in a stable probabilistic environment using a large adolescent and young adult sample with continuous age in the 8-30 range. Combining behavioral analysis and computational modeling, we showed developmental gains in performance from age 8-25 that were explained by an increase in learning from rewarded outcomes (corresponding to a narrower time window of information integration) and a decrease in exploration. These data and models help explain why learning and decision making differ during development and why a ‘one-size-fits-all’ approach may not equally serve youth at different stages.

## Supporting information

### Exclusion criteria details

We collected data from 297 participants on the Butterfly task. After an initial exclusion of 21 participants who were more likely to switch than stay after positive feedback, 276 participants remained. To further identify participants who were not engaged in the task without excluding participants solely on a pure performance criterion, we instead implemented the following, less stringent, conjunctive exclusion criteria:

- Criterion 1. Proportion of stay trials (the participant picked the same flower as the previous trial regardless of the butterfly) was higher than *median* + 2 * *sd*, across the whole group.
- Criterion 2. Proportion of stay trials was lower than *median* − 2 * *sd*, across the whole group.
- Criterion 3. The number of contiguous stay trials was higher than 12 contiguous stay trials in a row.
- Criterion 4. Any number of missing trials: available data less than the full 120 trials, indicating that participants stopped before the end of the experiment.
- Criterion 5. Performance was not better than chance (50%): performance was based on proportion of trials where participants correctly chose the preferred flower of the butterfly.

We excluded participants who fit both Criterion 5 and one of Criterion 1-4. S1 Table shows the detail breakdown of participants’ age group for each criterion.

**S1 Table.** Number of participants excluded due to each exclusion criterion for each of the four age groups.

| Age group | 8 - 13 | 13 - 18 | 18 - 25 | 25 - 30 |
| --- | --- | --- | --- | --- |
| Criterion 1 | 3 | 2 | 1 | 0 |
| Criterion 2 | 4 | 0 | 1 | 0 |
| Criterion 3 | 6 | 4 | 2 | 4 |
| Criterion 4 | 6 | 1 | 0 | 1 |
| Criterion 5 | 14 | 5 | 2 | 2 |
| Criterion 1-4 | 14 | 5 | 3 | 5 |
| Criterion 5 and 1-4 | 8 | 2 | 0 | 1 |

After applying this conjunctive criteria, we further eliminated 11 participants for later analysis. 1 more subject, despite above chance performance, was excluded for only having 18 trials of data. In the end, we analyzed *N* = 264 subjects, with 157 subjects younger than 18. S1 Fig shows more detailed age group and sex break down. Note that the final criterion is less stringent than a pure performance criterion.

**S1 Fig. Demographics.**

### Model comparison extended

Although the *α*^+^*α*^−^*βf* model has better model comparison score, we decided to go with the simpler *α*^+^0*βf* model as the winning model because (1) *α*^−^ was not recoverable and (2) *α*^+^0*βf* validated participants’ behavior better.

For fitted parameters, it is important to first verify if we can recover them from simulated data [34]. Specifically, because RL models are generative, we can simulate artificial data from fitted parameters and fit the model on the artificial data again to check the identifiability of the model parameters and the robustness of the model fitting procedure.

For the *α*^+^*α*^−^*βf* model, we were unable to recover the *α*^−^ parameter (S2 Fig A). For the *α*^+^0*βf* model, all three parameters could be reasonably recovered (S2 Fig B-D).

**S2 Fig. Generate and recover.**

We further validated the fitted models by visualizing the learning curves of simulated data and comparing with participants’ data S3 Fig. The *α*^+^0*βf* tracks participants’ learning curve well throughout the experiment, whereas the *α*^+^*α*^−^*βf* overshot during the middle segment of the experiment. This combined with *α*^−^ not recoverable suggests that the *α*^+^*α*^−^*βf* model, while having a better model comparison score (Fig 2B), has the risk of overfitting.

**S3 Fig. Model validation.**

### Pubertal effects extended

Here we aggregate all results for pubertal effects on model parameters for participants under age 18. We binned participants differently based on age, PDS and T1. To control for age, participants were broken into 4 equal-sized groups within each sex respectively and then combined. For PDS, we consider all participants with PDS score of 1 as 1 group, and broke down the rest into 3 equal-sized bins within each sex and then combined. For T1, we first log-transformed raw testosterone levels, and then broke down participants into 4 equal-sized bins within each sex and then combined the bins across sex.

We found a marginally significant effect of PDS on *α*^+^, but not on *β* or the forgetting parameter *f* (2-way ANOVA, *α*^+^: *F* (3, 148) = 0.05; *β*: *F* (3, 148) = 0.19; *f* : *F* (3, 148) = 0.7). There was no significant effect of sex or interaction with sex (2-way ANOVA, all *p*’s > 0.17).

We found a main effect of T1 on *α*^+^, but not on *β* or *f* (2-way ANOVA, *α*^+^: *F* (3, 141) = 0.001; *β*: *F* (3, 141) = 0.1; *f* : *F* (3, 141) = 0.27). There was again no effect of sex or interaction with sex (2-way ANOVA, all *p*’s > 0.4).

To test pubertal effects on model parameters in addition to age, within each age bin, we used either PDS or T1 to predict model parameters with linear regression. After controlling for age (S4 Fig), we did not find significant effects of PDS on *α*^+^ in any of the 4 age bins (linear regression, all *p*’s > 0.37). We did find a significant effect of T1 on *α*^+^ (S5 Fig) in the third age bin (linear regression, *β*_*T*1_ = 0.0008, *p* = 0.005), but not in the other 3 age bins (linear regression, all *p*’s > 0.17).

**S4 Fig. PDS effect on *α*^+^ in each of the four age groups younger than age 18.**

**S5 Fig. T1 effect on *α*^+^ in each of the four age groups younger than age 18.**

## Acknowledgments

We thank the many people who contributed to this project: Amy Zou, Lance Kriegsfeld, Celia Ford, Jennifer Pfeifer, Megan Johnson, Vy Pham, Rachel Arsenault, Josephine Christon, Shoshana Edelman, Lucy Eletel, Neta Gotlieb, Haley Keglovits, Julie Liu, Justin Morillo, Nithya Rajakumar, Nick Spence, Tanya Smith, Benjamin Tang, Talia Welte, and Lucy Whitmore. We would also like to thank our participants and their families. The work was funded by National Science Foundation SL-CN grant 1640885 to RD, AGEC, and LW.

